# Highly efficient genome editing in *Bacillus subtilis* via miniature DNA nucleases IscB

**DOI:** 10.1101/2025.02.11.637773

**Authors:** Jie Gao, Hongjie Tang, Yuhan Yang, Hengyi Wang, Qi Li

## Abstract

Existing CRISPR-based genome editing tools are limited in *Bacillus subtilis* due to the large *cas* gene. The recently reported DNA nuclease IscB has the potential to be developed into a novel genome editing tool due to its size being one-third of Cas9, while its application in *B. subtilis* remains unexplored. In this study, genome editing tools pBsuIscB/pBsuenIscB based on IscB and enIscB (enhanced IscB) were established in *B. subtilis* SCK6, and successfully deleted 0.6 kb to 4.3 kb genes with efficiencies up to 100%. Subsequently, the pBsuenIscB with higher deletion efficiency was used, whereby the large genomic fragment of 37.7 kb or 169.9 kb was deleted with only one ωRNA. Additionally, single-copy or multi-copy *mCherry* genes was integrated by using pBsuenIscB. Finally, the editing plasmid was eliminated and the second round of genome editing was completed. Overall, this study has successfully applied IscB to *B.subtilis*, expanded the genome editing toolbox of *B. subtilis*, and will help to construct *B. subtilis* chassis for production of a variety of biomolecules.

## INTRODUCTION

*Bacillus subtilis* (*B. subtilis*) is a Gram-positive bacterium with strong protein secretion ability and no endotoxin^1^. It is a generally recognized as safe bacterium and is widely utilized in the industrial production of various chemical products and recombinant proteins^2, 3^. Genome editing tools are indispensable for both exploring the fundamentals of *B. subtilis* life sciences and applying them in industrial fermentation^4, 5^.

The RNA-guided DNA nuclease could cleave the genomic DNA under the guidance of guide RNA (gRNA) to generate a double-strand break (DSB) and the genome could be repaired by providing homologous arms, thus enabling precise genome editing^6^. Based on the DNA nucleases Cas9 and Cpf1, single-plasmid^7–9^ and dual-plasmids^9–13^ genome editing systems were established in *B. subtilis*. However, limitations were presented in these editing systems. For example, a single gRNA only enabled genome editing of a single gene or short DNA fragment. In contrast, deletion of the large genomic fragments depended on two gRNAs, and the construction of these gRNAs was time-consuming and laborious^12^. In addition, the single plasmid system was inefficient in delivery due to the long length of the genes encoding Cas and the repair templates, which increased the difficulty of genome editing. Although the dual-plasmids system separated the gene encoding Cas and the gRNA with repair template onto two plasmids, the multi-plasmids transformation was complex and required more sophisticated antibiotics for plasmids stability. To address the above limitations, the development of miniaturized gene editors was considered as a novel and alternative strategy.

Recent studies reported miniature RNA-guided DNA nuclease IscB, encoded by the IS200/IS605 superfamily transposon, which was hypothesized to be the ancestor of Cas9 and was only one-third the size of Cas9^14^. Similar to the Cas9 protein, the IscB protein also contained HNH and RuvC domains and could form a ribonucleoprotein complex after binding with OMEGA RNA(ωRNA). This complex could recognize the target adjacent motif (TAM) sequence (5’-CAGGAA-3’) and the target sequence (16 bp) to cleave the DNA^15^. IscB was developed as a base editor for successful application in mammals, and the enIscB (enhanced IscB) system with higher base editing activity was further established in mammals^16^. Additionally, IscB has been modified by other researchers to establish highly active miniature base editors in human and mouse cells respectively^17, 18^. Besides, a novel IscB was mined from the macro-genome database and developed IscB.m16*-BE, a miniature base editing tool with a wider TAM sequence was extended from MRNRAA to NNNGNA^19^.

The above reports elaborated that the miniature DNA nuclease IscB had great potential in genome editing applications, but the existing related studies on IscB mainly focused on genome editing in eukaryotes, the application of IscB in prokaryotes(eg *B. subtilis*) had not been reported. Therefore, *B. subtilis* SCK6 was selected as the host to test the application of IscB in this study. Firstly, the toxicity performance and the ability to cleave genomic DNA of IscB and enIscB in *B. subtilis* SCK6 were verified. Then, genome editing systems pBsuIscB/pBsuenIscB based on IscB/enIscB were established, in which the pBsuenIscB system showed a slightly higher editing efficiency, so it was used in the following researches. Subsequently, the large genomic fragments with lengths of 37.7 kb or 169.9 kb was successfully deleted by using only a single ωRNA. In addition, single-copy or multi-copy *mCherry* genes was integrated into the *B. subtilis* SCK6 respectively. Finally, the pBsuenIscB system was eliminated, and the second round of genome editing was successfully completed. In this study, a genome editing tool based on IscB was successfully established in *B. subtilis* for the first time, which expanded the genome editing toolbox of *B. subtilis* and will accelerate the construction of *B. subtilis* cell factories.

## MATERIALS AND METHODS

### Strains and Culture Conditions

The strains used in this study were listed in Table S1. *Escherichia coli* (*E. coli*) DH5α was used as the cloning host for plasmid construction, *B.subtilis* SCK6 served as the genome editing host. *E.coli* and *B. subtilis* SCK6 were cultivated in LB medium (0.5% [w/v] yeast extract, 1% [w/v] tryptone, and 1% [w/v] NaCl) at 37°C, 220 rpm. In LB medium, 50 μg/mL and 20 μg/mL kanamycin were added to screen plasmids of *E. coli* and *B. subtilis* SCK6, respectively. Competent cells of *B. subtilis* SCK6 were prepared using YN medium that contained 0.7% [w/v] yeast extract, 1% [w/v] tryptone, 0.5% [w/v] NaCl, and 0.3% [w/v] beef extract. Additionally, 1.5% [w/v] xylose was added to induce the expression of the *comk* gene to form competent cells of *B. subtilis* SCK6. Strains were stored at -80°C at a final concentration of 20% glycerol, and the strain recovery was carried out by streaking the culture on LB plate and inoculating the isolated colony into LB broth. Add uracil (10 mg/L) as required.

### Reagents and Enzymes

The restriction enzymes applied in this study were purchased from Thermo Fisher Scientific (Waltham, MA, USA). DNA polymerase 2×Phanta Flash Master Mix (Vazyme Biotechnology Co., Ltd., Nanjing, China) and 2×Es Taq Master Mix (Dye) (Jiangsu Cowin Biotech Co., Ltd., Taizhou, China) were used for high-fidelity DNA amplification and *E. coli* colony PCR respectively. 2×Rapid Taq Master Mix (Vazyme Biotechnology Co., Ltd., Nanjing, China) were used for *B. subtilis* SCK6 colony PCR. The reagent kit used for plasmid extraction and DNA purification was obtained from Tiangen (Tiangen Biotech Co., Ltd., Beijing, China), according the manufacturer’s instructions. The ClonExpress One Step Cloning Kit (Vazyme427 Biotechnology Co., Ltd., Nanjing, China) was used for the assembly of plasmids.

### Plasmids Construction

All plasmids and primers used in this study were listed in supplementary tables S1 and S2. *E.coli* DH5α was used for plasmid maintenance and construction. The IscB and enIscB gene sequences and corresponding ωRNA were respectively derived from plasmids IscB-ωRNA and enIscB-ωRNA. Taking the IscB-related plasmids as an example, the xylose induced promoter P*_xylA_* (expressing IscB) was amplified from pHT43-XCR6^11^ and assembled with the IscB gene sequence into the linearized pcrF19-NM2^11^ plasmid vector, and the pUC18 replicon was replaced with the p15A replicon with a lower copy number to generate the pIscB plasmid. The ωRNA and its constitutive promoter P*_veg_* were superimposed on the pIscB plasmid to obtain the cleavage plasmid pBsuIscB-ωRNA-targetX (targetX represents the target gene).

For the pBsuIscB-targetX plasmid, it could be obtained by inserting the repair template (1 kb upstream and 1 kb downstream of the homologous arms) amplified from the genome of *B. subtilis* SCK6 into pBsuIscB-ωRNA-targetX. The construction of enIscB-related plasmids was similar to the above method. For the integration plasmid, the integration gene and its promoter were added into the upstream and downstream homologous arms of the target gene.

### Preparation of Competent Cells of *B. subtilis* SCK6 and Plasmid Transformation

The *B. subtilis* SCK6 stored at -80℃ was streaked on LB plates for activation. Subsequently, the single colonies obtained from streaking were inoculated into fresh LB medium. The culture was incubated at 37℃, 220 rpm for 12 h. Then the overnight culture was transferred to fresh YN medium containing 1.5% xylose, with an initial OD_600_ of 1.0, and further incubated at 37℃, 220 rpm for 2 h to form competent cells.

Plasmids were mixed with 500 µL of competent cells, and the mixture was incubated at 37°C, 220 rpm, 90 min and then spread on LB plates.

### The Tests of Toxicity and Genome Cleavage

We added 5 µL of pIscB/penIscB plasmid to 500 µL of *B.subtilis* SCK6 competent cells, the mixture was spread on LB plates with 3% xylose and kanamycin (experimental group) and LB plates with only kanamycin (control group) after transformation, and incubated overnight at 37°C. Colony-forming units (CFU) were counted to calculate the plasmid transformation efficiency (CFU/µg). If the transformation efficiency of the experimental group was significantly lower than that of the control group, it was considered that the IscB/enIscB was toxic to *B.subtilis* SCK6. Conversely, if the transformation efficiency of the experimental group was equivalent to that of the control group, it was considered that the protein was not toxic to *B.subtilis* SCK6, and the cutting test of the corresponding protein could be continued.

For the test of genome cleavage, the steps were similar to the above method. Plasmids were pBsuIscB-ωRNA-targetX and pBsuenIscB-ωRNA-targetX. If the transformation efficiency of the experimental group on the LB plate was similar to that of the control group, it was considered that the protein could not cleave the host genome DNA. Conversely, if the transformation efficiency of the experimental group was significantly lower than that of the control group, it was considered that the protein could cleave the genome DNA of *B.subtilis* SCK6.

### Gene deletion and gene insertion

We added 10 µL pBsuIscB-targetX/pBsuIscB-targetX plasmid with a known concentration to 500 µL of *B. subtilis* SCK6 competent cells, mixed well, and then incubated for 90 min at 37°C, 220 rpm, after which they were spread on LB plates containing 3% xylose and kanamycin (20 μg/mL) and cultured at 37℃ overnight. Fifteen transformants were randomly selected on selected plates for PCR. The deletion primers complementary to the upstream ∼50 bp and downstream ∼50 bp of the homology arms on the genome were used. The insertion primers were respectively located within the integration gene and ∼50 bp outside the downstream homologous arm. The PCR products were subsequently sequenced to verify the positive colonies, and the wild-type strain were used as control. For the integration of multi-copy *mCherry* genes, quantitative PCR (qPCR) was used to verify the copy number of the *mCherry* genes on the genome. The *pyrE* deficiency conferred a uracil auxotrophic phenotype on *B. subtilis*, and the screening plates for strains with the *pyrE* gene deleted required the addition of uracil (10 mg/L).

### Determine the copy number of the *mCherry* genes by qPCR

The strain with the single-copy *mCherry* gene integrated at the *spo0A* site was used as the standard sample. The genomic DNAs (gDNA) of the standard sample and the strains with *mCherry* gene integrated at the *rrn* operon were extracted and diluted as the template. The *dnaN* gene was choesn as the single-copy reference gene on the genome of *B. subtilis* SCK6. Specific primer pairs dnaN qup/dnaN qdn and mCherry qup/mCherry qdn (Table S2) were designed to generate products of approximately 180 bp. The PCR mixture (20 μL) contained 10 μL of 2×Taq Pro Universal SYBR qPCR Master Mix, 0.4 μL of L forward and reverse primers (10 μM), 10 ng genome DNA, and H2O. The reaction condition was as followed: 95℃ for 30 s; 40 cycles of 95℃ for 10 s, 60℃ for 10 s; melt curve 65°C to 95℃ in 0.5°C/5 s increments. Copy-numbers for all samples were calculated by Pfaffl Method. All experiments were performed with three independent biological replicates.

### Plasmid Curing

To eliminate the editing plasmid pBsuenIscB, the colony of the edited clone containing pBsuenIscB was inoculated into 5 mL of LB medium supplemented with kanamycin (20 μg/mL). The culture was incubated overnight with shaking at 220 rpm. Approximately 10 μL of the cells were transferred into antibiotic-free LB and incubated at 50°C with 200 rpm for 8 h. The incubation was continued three times at 50°C with 200 rpm for 8 h, and then diluted and spread on antibiotic-free LB plates and incubated overnight at 37°C. Colonies growing on the LB plates were randomly picked and screened by spotting on LB plates with kanamycin (20 µg/mL) and without kanamycin. The colonies that were sensitive to kanamycin were cured of pBsuenIscB. The same method could be used for the elimination of pBsuIscB.

## RESULTS AND DISSCUSION

The genome cleavage of *B. subtilis* SCK6 was achieved by using IscB.

The recently reported RNA-guided DNA nuclease IscB poses the advantage of a small size (496 amino acids), and it has been shown to work in mammalian cells^17, 18, 20^. Some researchers have engineered IscB and its corresponding ωRNA derived from the human intestinal macrogenome to obtain an enhanced IscB version named enIscB, which further improves editing activity^16^. However, studies of IscB and enIscB in bacteria have not been reported. Therefore, this study aimed to explore whether IscB and enIscB could be utilized for genome editing in *B. subtilis* SCK6. Successful genome editing requires that IscB should be nontoxic to *B. subtilis* SCK6. Thus, plasmids pIscB (encoding IscB, expressed under the P*_xylA_* inducible promoter) and penIscB (encoding enIscB, expressed under the P*_xylA_* inducible promoter) were respectively transformed into *B. subtilis* SCK6, and the transformants were spread on LB plates (containing 3% xylose and kanamycin) and LB plates (containing only kanamycin). Then, the number of surviving colonies on the plates was counted and the transformation efficiency was calculated (Fig 1A). The results showed that both pIscB and penIscB could grow on LB plates with or without xylose, and their transformation efficiencies were comparable, indicating that the IscB and enIscB proteins were non-toxic to *B. subtilis* SCK6 (Fig 1B and Fig S1). Moreover, the transformation efficiency of penIscB was slightly higher than that of pIscB.

**Fig 1.**
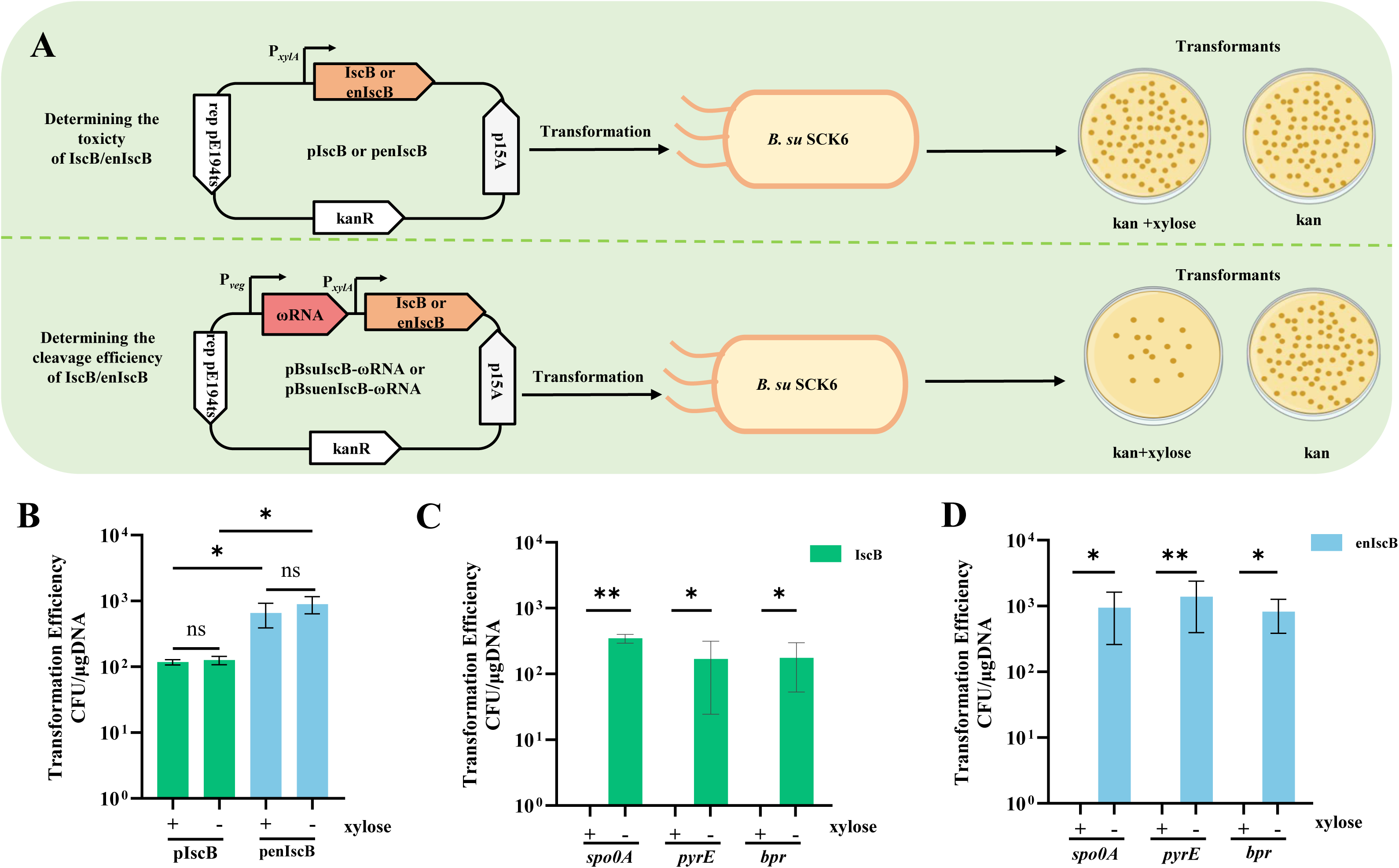
IscB could be utilized to achieve the genome cleavage of *B. subtilis* SCK6. (A) The toxicity tests of pIscB/penIscB in *B. subtilis* SCK6 (above) and the tests on the cleavage of the genome of *B. subtilis* SCK6 by pBsuIscB-ωRNA/penIscB-ωRNA (below). +xylose: the LB plate with 3% xylose to induce the expression of IscB/enIscB; -xylose: the LB plate without 3% xylose, and IscB/enIscB was not expressed. (B) The toxicity results of pIscB/penIscB in *B. subtilis* SCK6. (C) The efficiency of the genome cleavage by using pIscB. (D) The efficiency of the genome cleavage by using penIscB. ns, not significant; *p < 0.05; **P < 0.01.

Next, the genome cleavage function of IscB and enIscB in *B. subtilis* SCK6 was tested. The ωRNA was inserted into the pIscB and penIscB plasmids to obtain cleavage plasmids named pBsuIscB-ωRNA and pBsuenIscB-ωRNA, respectively. The cleavage plasmids pBsuIscB-ωRNA (MolecularCloud number: MC_0101504) and pBsuenIscB-ωRNA (MolecularCloud number: MC_0101505) carrying ωRNA that target *pyrE*, *spo0A*, or *bpr* gene were transformed into *B. subtilis* SCK6, respectively. The transformants of experimental groups (cleaved genome) were spread on LB plates containing 3% xylose and kanamycin, and the control groups transformants (un-cleaved genome) were spread on LB plates containing only kanamycin (Fig 1A). Notably, the transformation efficiency of the experimental groups was nearly 0 CFU/μg DNA, and that of the control groups was 10^3^ CFU/μg DNA (Fig 1C-D and Fig S2-3). It was demonstrated that both IscB and enIscB proteins were able to cleave the genome of *B. subtilis* SCK6.

These above findings revealed that IscB and enIscB were non-toxic in *B. subtilis* SCK6 and could also cleave the genome with high efficiency. Thus, it was implied that it was promising to further achieve genome editing in this host by providing a repair template with IscB and enIscB^16^. In addition, the TAM sequences of IscB and enIscB (5’-CAGGAA-3’) are longer than the PAM sequence of Cas9 (5’-NGG-3’) and Cpf1 (5’-TTTN-3’). Due to the longer TAM sequences, IscB and enIscB can localize the target gene locus more accurately and reduce the incorrect binding with non-target sequences. Consequently, they demonstrate higher specificity in identifying target sequences and are expected to decrease the off-target rate, thus having favorable application prospects in genetic engineering research and related application fields.

### Establishing the IscB-based genome editing system in *B. subtilis* SCK6

To validate the genome editing efficiency of IscB and enIscB in *B. subtilis* SCK6, *pyrE*, *spo0A* and *bpr* genes were selected for the testing of deletion. The upstream (∼1 kb) and downstream (∼1 kb) homology arms of the deletion gene were added on pBsuIscB-ωRNA and pBsuenIscB-ωRNA to obtain genome editing systems named pBsuIscB (MolecularCloud number: MC_0101506) and pBsuenIscB (MolecularCloud number: MC_0101507). Then, the genome editing systems were transformed into *B. subtilis* SCK6, and 15 transformants were randomly selected for checking the gene deletion (Fig 2A). Corresponding TAM and N16 sequence information for *pyrE*, *spo0A* and *bpr* genes was shown in Fig 2B. The results showed that the gene deletion efficiencies of the pBsuIscB system were 93.33% (*pyrE*), 13.33% (*spo0A*), and 100% (*bpr*), while those of the pBsuenIscB system were 26.67% (*pyrE*), 100% (*spo0A*), and 100% (*bpr*) (Fig 2C). The mutants were further verified by DNA sequencing to confirm that the above genes were successfully deleted (Fig 2D-G).

**Fig 2.**
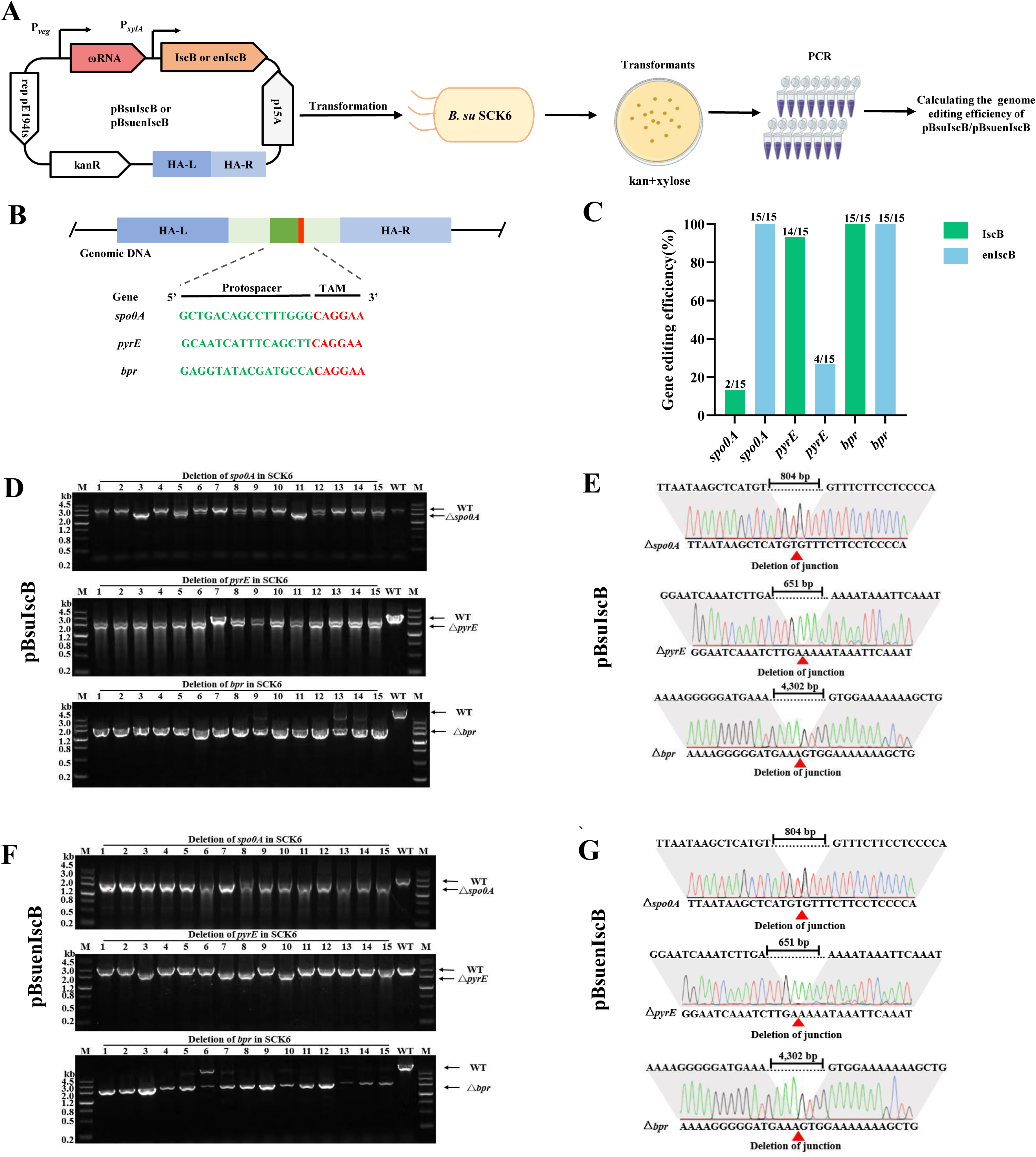
The IscB-based genome editing system was established in *B. subtilis* SCK6. (A) Schematic representation of gene deletion in *B. subtilis* SCK6 based on the pBsuIscB/pBsuenIscB system. (B) The target sequence of different target genes in *B. subtilis* SCK6. (C) Gene deletion efficiency of the pBsuIscB/pBsuenIscB system. (D) DNA gel images for PCR verification of the deletion of *spo0A*, *pyrE* and *bpr* genes by the pBsuIscB system. (E) DNA sequencing results of the deletion genes by the pBsuIscB system. Lengths of the deleted genes: Δ*spo0A*: 804 bp, Δ*pyrE*: 651 bp, Δ *bpr*: 4302 bp. (F) DNA gel images for PCR verification of the deletion of *spo0A*, *pyrE* and *bpr* genes by the pBsuIscB system. (G) DNA sequencing results of the deletion genes by the pBsuenIscB system. Lengths of the deleted genes: Δ*spo0A*: 804 bp, Δ *pyrE*: 651 bp, Δ*bpr*: 4302 bp.

These above results demonstrated that the gene deletion in *B. subtilis* SCK6 could be achieved by using pBsuIscB and pBsuenIscB systems, and the efficiency could reach as high as 100%, which was equivalent to the gene deletion efficiency of the CRISPR system^21^. Further analysis revealed that the pBsuenIscB system had a better performance in editing efficiency compared with the pBsuIscB system. We speculated that this might be related to the fact that enIscB were engineered from IscB, since the editing activity of enIscB was indeed higher than that of IscB in mammals^16^. Since the pBsuenIscB system had a more effective editing efficiency, it was employed for the further applications.

### Deletion of the large genomic fragments can be achieved in *B. subtilis* SCK6 by p BsuenIscB system

Although gene deletions in *B. subtilis* SCK6 were achieved above, the size of the genes tested was not large. In practice, to improve the efficiency of microbial cell factories, it usually involves deleting non-essential regions to reduce transcription costs and eliminate competing pathways, and these non-essential regions are tens of kilobases or even more than 100 kb in size^22–24^. Currently, only one method for deletion of large genomic fragments was established in *B. subtilis* by CRISPR-Cas9^12^. However, it requires the design of two gRNAs at both ends of the large genomic fragment respectively to generate DSBs, which makes the operation cumbersome.

To evaluate the potential of the pBsuenIscB system for deleting large genomic fragments in *B. subtilis* SCK6, the *pps* operon (non-essential for cell growth) with a length of 37.7 kb was chosen as the target. The homology arm (∼1 kb for upstream and ∼1 kb for downstream) and the single ωRNA targeting the middle of the *pps* operon were added to penIscB to yield the editing plasmid pBsuenIscB-*pps* (Fig 3A). Then, the editing plasmid pBsuenIscB-*pps* was transformed into *B. subtilis* SCK6, and 15 transformants were randomly selected for PCR verification. Since the primers were designed outside the homology arm on the genome, thus, the 2 kb band remained because of the deletion of the *pps* operon. In contrast, the band of the wild-type strain was approximately 40 kb and no results appeared. Fortunately, the *pps* operon was successfully deleted with an efficiency of 100% (Fig 3B). In order to further verify whether the larger genomic fragments could be deleted by using pBsuenIscB system along with a single ωRNA, a 169.9 kb genomic fragment was selected for deletion (Fig 3C). Surprisingly, this work was also successfully completed by the pBsuenIscB system, with an efficiency of 40% (Fig 3D).

**Fig 3.**
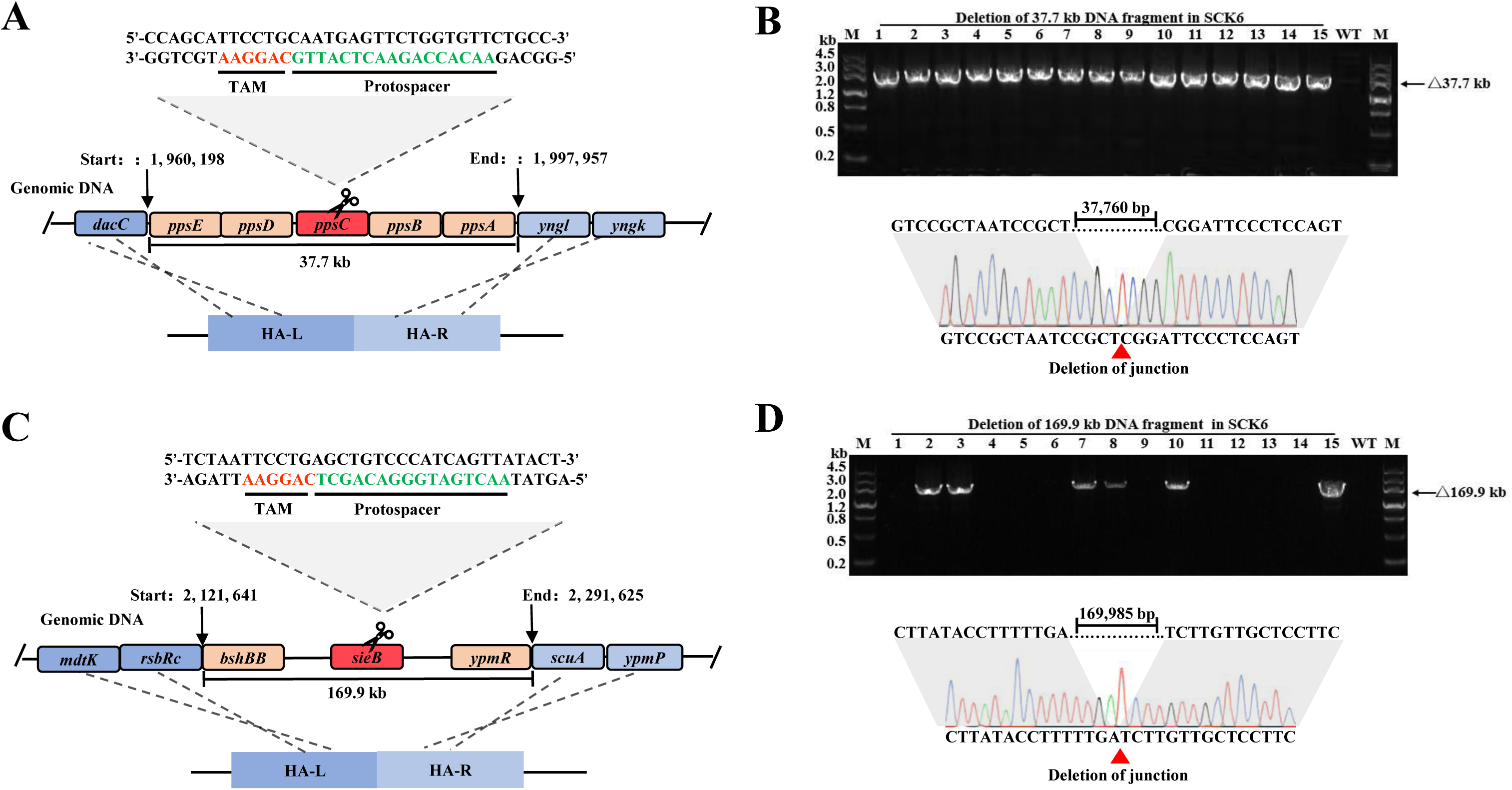
The pBsuenIscB system enabled the deletion of large genomic fragment in *B. subtilis* SCK6. (A) Schematic diagram of the target site information and homologous arms for the deletion of the 37.7 kb *pps* operon. (B) DNA gel image and sequencing results of the deletion of the *pps* operon by the pBsuenIscB system. (C) Schematic diagram of the target site information and homologous arms for the deletion of the 169.9 kb large genomic fragment. (D) DNA gel image and sequencing results of the deletion of the 169.9 kb large genomic fragment.

Here, the pBsuenIscB system accomplished the deletion of large genomic fragments of 37.7 kb and 169.9 kb. Notably, our system only required one ωRNA to delete large genomic fragments, while it required two gRNAs for CRISPR-Cas9 system^12^. A single ωRNA avoided the complexity of combining multiple elements, simplified the process of plasmid construction made the operation process more convenient and reduced the time consumption.

### Gene integration can be achieved in *B. subtilis* SCK6 by pBsuenIscB system

Currently, plasmids are mainly used in industrial production to increase the synthesis of target products. However, plasmids need to rely on antibiotics to be maintained and are unstable in host bacteria. To achieve the stable expression of genes, the gene expression cassette can be integrated into the genome of the host^25^. To examine the efficiency of the pBsuenIscB system in gene integration, the *mCherry* gene that expresses the red fluorescent protein was used as a reporter gene and integrated into the *spo0A* site of *B. subtilis* SCK6. The integrating plasmid pBsuenIscB-*spo0A*-*mCherry* was obtained by inserting the *mCherry* gene (controlled by P*_spl_* promoter) between the upstream and downstream homologous arms of pBsuenIscB-*spo0A* (Fig 4A), and then transformed into *B. subtilis* SCK6 for gene integration. By randomly selecting 15 transformants for PCR, it was verified that the pBsuenIscB system could successfully achieve *mCherry* gene integration with efficiency of 20% (Fig 4B and Fig S4). The obtained single-copy *mCherry* gene strain was named Mut-SC-*mCherry*.

**Fig 4.**
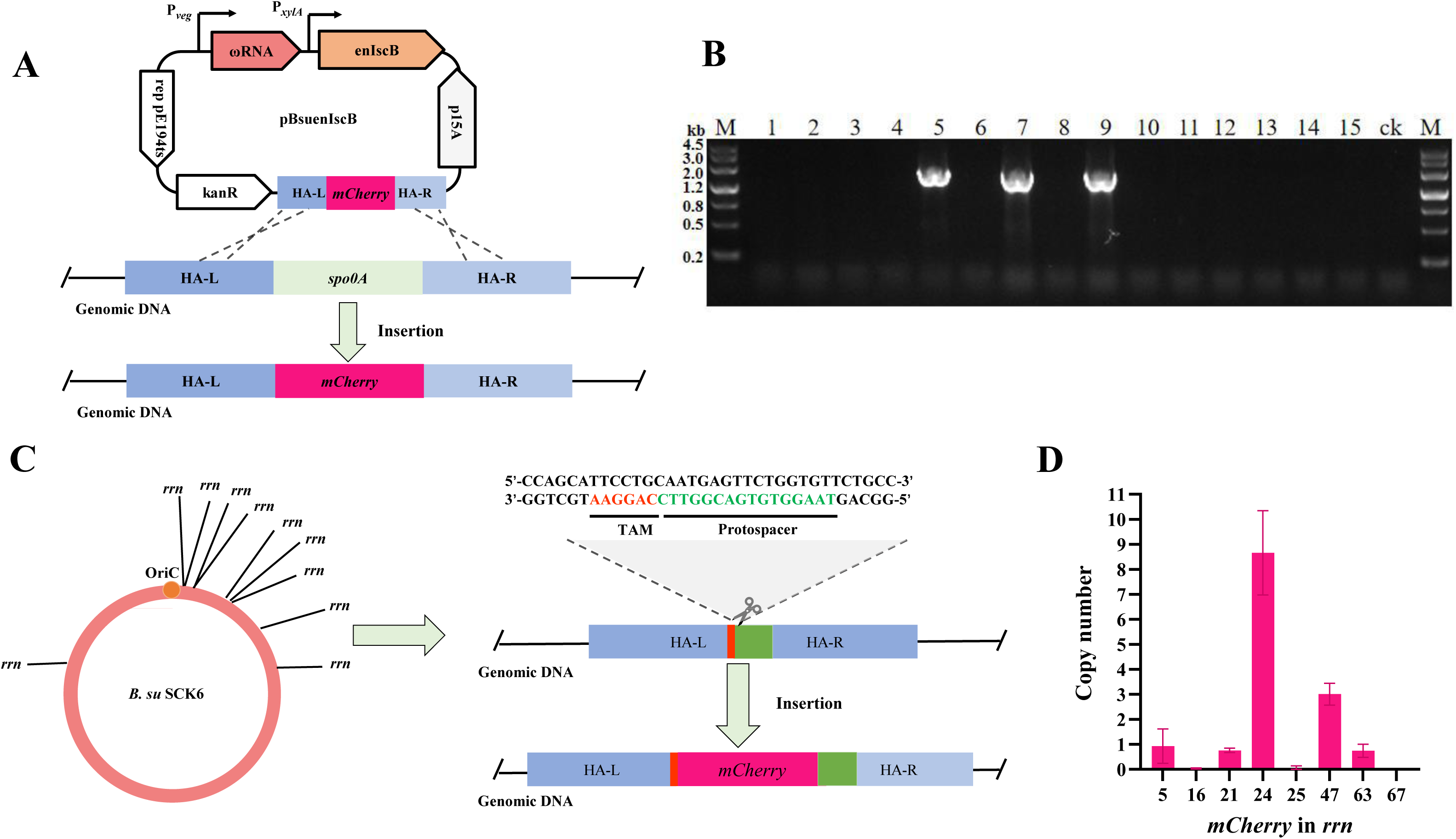
Single-copy and multi-copy *mCherry* genes were integrated in *B. subtilis* SCK6 by pBsuenIscB system. (A) Schematic diagram of the integration of the single-copy *mCherry* gene at the *spo0A* site. (B) DNA gel image of the integration of the single-copy *mCherry* gene. (C) Distribution of the *rrn* operon in the genome of *B. subtilis* SCK6 and schematic diagram of the integration of the multi-copy *mcherry* genes. (D) The copy number of the *mCherry* gene in the integrated strains.

Previous studies found that there were 10 copies of the ribosomal RNA (*rrn*) operon (encoding bacterial ribosomal 23S, 16S and 5S rRNAs) in the genome of *B. subtilis*, and damage to them would affect the growth rate of *B. subtilis*. The effect on the growth rate of *B. subtilis* was significant when the copy number of *rrn* operon was less than two, while the growth rate at the remaining copy numbers was acceptable^26^. It was verified that integrating the target gene at the *rrn* operon was feasible. To further increase the expression level of the *mCherry* gene, an attempt was made to integrate multiple copies of the *mCherry* gene into the *rrn* operon (Fig 4C). The integrating plasmid pBsuenIscB-*rrn*-*mCherry* was transformed into *B. subtilis* SCK6. Then, a total of 69 transformants were picked to screen for positive colonies, with an integration efficiency of 39.7% (Fig S5 A-B). The multi-copy *mCherry* genes strain was named Mut-MC-*mCherry*. Subsequently, genomic DNA (gDNA) of eight Mut-MC*-mCherry* strains was extracted for qPCR to confirm the copy number of the *mCherry* gene, with the gDNA of Mut-SC-*mCherry* as the standard sample and the *dnaN* gene as the reference gene. Fortunately, strains 24 and 47 were integrated *mCherry* genes with copy numbers 8 and 3 respectively, and the rest were single-copy strains (Fig 4D).

Previously, there were mainly two methods for the multi-copy integration research on *B. subtilis*. The first method was to integrate single genes one by one^27^, which was time-consuming, laborious and had low efficiency. The second method was to first integrate the *crtMN* operon (encoding xanthophyll) into three sites of the *B. subtilis* genome, and then with the help of the CRISPR-Cas9 system, it was possible to integrate up to three copies of exogenous genes into *B. subtilis* simultaneously^28^. In this study, a more subtle strategy of targeting 10 copies of *rrn* operon simultaneously with only one ωRNA was used to randomly achieve the integration of 3 and 8 copies of *mCherry* genes in *B. subtilis* SCK6, which further enhanced the integration rate of multicopy genes.

This research fully demonstrated that the pBsuenIscB system had the ability to achieve single-copy *mCherry* gene integration at the *spo0A* site and multi-copy *mCherry* genes integration at the *rrn* site, and this method could be used for the stable expression of exogenous target genes in the future.

### The curing of pBsuenIscB system and second round of genome editing

Plasmid curing is the prerequisite for testing phenotypes and second round of genome editing^29^. For the plasmid elimination test, pBsuenIscB-*bpr* was selected as the target. The correctly edited *B. subtilis* SCK6 Δ *bpr* strain was inoculated into antibiotic-free LB medium and then incubated at 50°C with shaking at 200 rpm for 8 h. This process of incubation and subculturing was repeated for 3 times. Subsequently, the bacterial solution was taken and diluted for spreading to obtain single colonies, followed by a kanamycin spotting test on these single colonies (Fig 5A). A total of 52 colonies were picked, among which 48 colonies were sensitive to kanamycin, suggesting that the editing plasmid pBsuenIscB-*bpr* had been cured with efficiency of 92.3% (Fig 5B).

**Fig 5.**
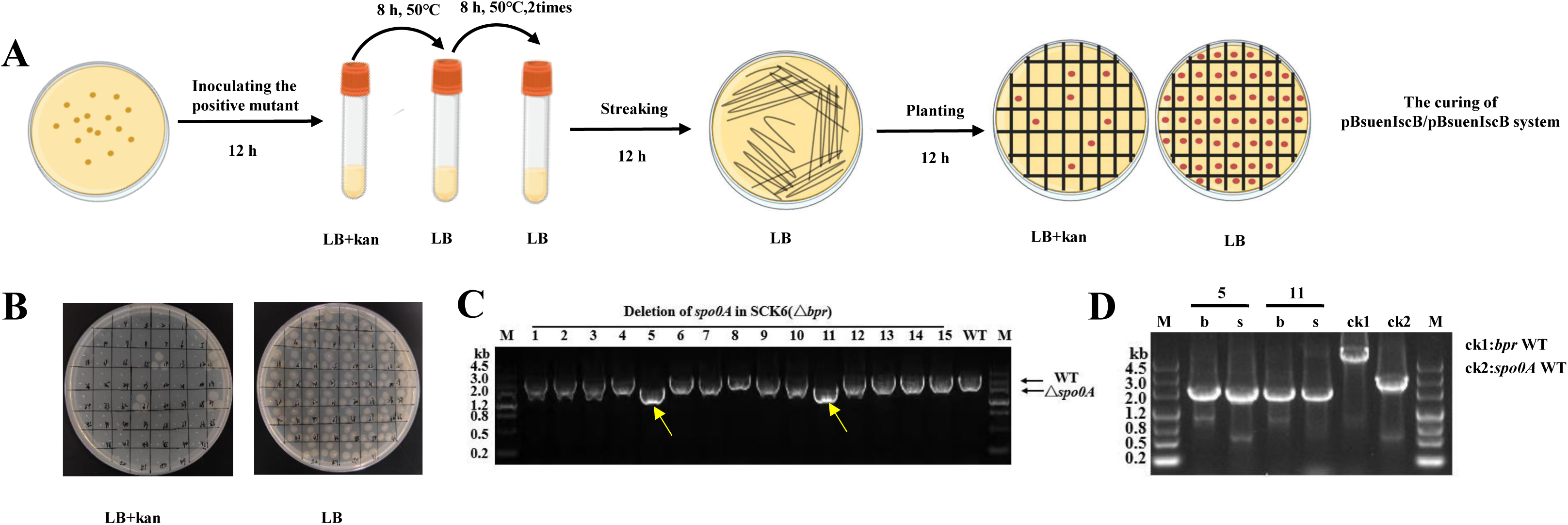
The pBsuenIscB system was cured and the second round of genome editing was accomplished successfully. (A) Operational procedures for the elimination of the pBsuenIscB system. (B) Resistance sensitivity spotting test to verify that the editing plasmid pBsuenIscB-*bpr* in *B. subtilis* SCK6 Δ *bpr* strain has been eliminated. (C) DNA gel image of the second-round genome editing at the *spo0A* site by using the pBsuenIscB system, strains No. 5 and No. 11 were the successfully edited strains. (D) Verification results of the deletion of the *bpr* and *spo0A* genes in strains No. 5 and No. 11.

For second round of genome editing, the editing plasmid pBsuenIscB-*spo0A* was transformed into the above *B. subtilis* SCK6 Δ *bpr* strain with the plasmid already cured. The results showed that second round of genome editing at the *spo0A* site was completed by using pBsuenIscB system with an efficiency of 13.3% (Fig 5C), and the deletions of *bpr* and *spo0A* genes were further verified by PCR (Fig 5D).

This research successfully cured the editing plasmid pBsuenIscB-*bpr* and achieved second round of genome editing using the pBsuenIscB system. It verified the feasibility of system for multi-target genomic editing and provides a basis for modifying engineered strains with the pBsuenIscB system.

## Acknowledgment

This study was supported by National Natural Science Foundation of China (32402894) and Sichuan Science and Technology Program (Number: 2024NSFSC0373). We would like to thank Prof. Hui Yang and Yingsi Zhou for providing the pIscB/penIscB-ωRNA related plasmids; Prof. Long Liu for providing the pHT-XCR6 and pcrF19-NM2 plasmids.

## CONFLICTS OF INTEREST

The authors declare no conflict of interest.

## Supplemental material

Strains, plasmids and primers used in this study; the results of the toxicity and the ability to cleave genomic DNA by IscB and enIscB in *B. subtilis* SCK6; sequencing results of single-copy and multi-copy *mCherry* genes integration(PDF).

## Notes

### Competing Interest Statement

The authors have declared no competing interest.

