## Supplemental information for "Highly efficient genome editing in *Bacillus subtilis* via miniature DNA nucleases IscB"

**Supplemental material.**

Table S1: Plasmids and strains used in this study.

| Strains or plasmids | Description | Source |
|---|---|---|
| Strains | | |
| *E. coli* DH5α | Commercial transformation host | GIBCO BRL, life Technologies |
| *B. su* SCK6 | Host for testing the genome editing efficiency | Lab storage |
| Plasmids | | |
| pHT-XCR6 | | (1) |
| pcrF19-$NM_2$ | | (1) |
| pIscB | Derived from pcrF19-NM2, inducibly expressing IscB | This study |
| penIscB | Derived from pcrF19-NM2, inducibly expressing enIscB | This study |
| pBsuIscB-ωRNA-*spo0A* | Derived from pIscB , target *spo0A* gene | This study |
| pBsuIscB-ωRNA-*pyrE* | Derived from pIscB, target *pyrE* gene | This study |
| pBsuIscB-ωRNA-*bpr* | Derived from pIscB, target *bpr* gene | This study |
| pBsuIscB-*spo0A* | Derived from pBsuIscB-ωRNA-*spo0A*, carry the homology arms of *spo0A* | This study |
| pBsuIscB--*pyrE* | Derived from pBsuIscB-ωRNA-*pyrE* , carry the homology arms of *pyrE* | This study |
| pBsuIscB-*bpr* | Derived from pBsuIscB-ωRNA-*bpr* , carry the homology arms of *bpr* | This study |
| pBsuenIscB-ωRNA-*spo0A* | Derived from penIscB , target *spo0A* gene | This study |
| pBsuenIscB-ωRNA-*pyrE* | Derived from penIscB , target *pyrE* gene | This study |
| pBsuenIscB-ωRNA-*bpr* | Derived from penIscB , target *bpr* gene | This study |
| pBsuenIscB-*spo0A* | Derived from pBsenuIscB-ωRNA-*spo0A*, carry the homology arms of *spo0A* | This study |
| pBsuenIscB-*pyrE* | Derived from pBsenuIscB-ωRNA-*pyrE* , carry the homology arms of *pyrE* | This study |
| pBsuenIscB-*bpr* | Derived from pBsuIscB-ωRNA-*bpr* , carry the homology arms of *bpr* | This study |

| | | |
|---|---|---|
| pBsuenIscB-ωRNA-*pps* | Derived from penIscB , target *pps* operon | This study |
| pBsuenIscB-*pps* | Derived from pBsenuIscB-ωRNA-*pps* , carry the homology arms of *pps* | This study |
| pBsuenIscB-ωRNA-169.9 | Derived from penIscB , target 169.9 genomic fragment | This study |
| pBsuenIscB-169.9 | Derived from pBsenuIscB-ωRNA-169.9 , carry the homology arms of 169.9 genomic fragment | This study |
| pBsuenIscB-*spo0A-mCherry* | Derived from pBsuenIscB-*spo0A* , carry the *mCherry* gene and $P_{spl}$ | This study |
| pBsuenIscB-ωRNA-*rrn* | Derived from penIscB , target *rrn* operon | This study |
| pBsuenIscB-*rrn-mCherry* | Derived from pBsenuIscB-ωRNA-*rrn* , carry the homology arms of *rrn*, the *mCherry* gene and $P_{spl}$ | This study |

Table S2: Oligonucleotides used in this study.

| Oligos | Sequence (5’→3’) |
|---|---|
| pcrf-part-up | ggcctgcagatctacgtgtaaagagtcgacctgttacgaa |
| pcrf-part-dn | caatgtttttttagaaaatttagttacgcatcacacgcaaaaaggaaat |
| IscB-up | agggggaaatgggatccatgatggccgtggtgtacgtgat |
| IscB-dn | ttcgtaacaggtcgactctttacacgtagatctgcaggc |
| PxylA-up | taactaaattttctaaaaaaaaacattg |
| PxylA-dn | ctgatcacgtacaccacggccatcatggatcccatttccc |
| IscB-verf-up | acaatggacacctagggg |
| IscB-verf-dn | accatgaaaaagagcccg |
| IscB-ce-1 | tgagcagccctaaaggcaga |
| IscB-ce2 | cgaagaagaagagccaagg |
| p15A-up | cgttccactgagcgtcagaccctagcggagtgtatactgg |
| p15A-dn | aacgcaggaaagaaattaataagatgatcttcttgagatc |
| p15A-part-up | tcaagaagatcatcttattaatttctttcctgcgttatcc |
| p15A-part-dn | taagccagtatacactccgctagggtctgacgctcagtgg |
| p15A verf up | taaagataccaggcgtttcc |
| p15A verf dn | tggaatgggattcaggagtg |
| ωRNA-dn | ctgggtttaagcttgcttgccgttcctccttttgtaaaac |

| | |
|---|---|
| ter-up | gttttacaaaaggaggaacggcaagcaagcttaaacccag |
| ter-dn | aatgttttttttagaaaatttagttagactctagaggcactggccaagc |
| pveg-dn | acatttattgtacaacacgagc |
| enIscB-ωRNA-spo0A | atgggctcgtgttgtacaataaatgtcgttgcttataacggaggctcgtccaactgcgg |
| IscB-ωRNA-spo0A | gctcgtgttgtacaataaatgtgctgacagcctttgggggctcttccaactttatggt |
| HR-part-spo0A-dn | cgggaaggctaatcgtctgagagatcccctcataatttca |
| HR-part-spo0A-up | tatcacgtttgatcagtaagtctgctcgaggtcatcgttc |
| spo0A-upup | tctcagacgattagccttcccgctc |
| spo0A-updn | ggaggaagaaacacatgagcttattaagtggtc |
| spo0A-dnup | gctcatgtgtttcttcctccccaaatgtag |
| spo0A-dndn | gcagacttactgatcaaacgtg |
| HR-verf-up | tttcggtttaccggtgtc |
| HR-verf-dn | tggaatgggattcaggagtg |
| spo0A-ko-up | cagcgcacggatcgctgg |
| spo0A-ko-dn | gccccatttattcaaaaggc |
| spo0A-ko-ce-up | gacacatctagggtcgatc |
| IscB-ωRNA-pyrE | ggctcgtgttgtacaataaatgtgcaatcatttcagcttggctcttccaactttatggt |
| enIscB-ωRNA-pyrE | gctcgtgttgtacaataaatgtgcaatcatttcagcttggctcgtccaactgcgg |
| HR-part-pyrE-dn | taatatcggtttgcctctgagagatcccctcataatttca |
| HR-part-pyrE-up | cgcttcatgaccaaggctatcgctcgaggtcatcgttcaa |
| pyrE-upup | ttatgaggggatctctcagaggcaaaccgatattagcg |
| pyrE-updn | acatcatttgaatttatttttcaagatttgattccctccc |
| pyrE-dnup | gggagggaatcaaatcttgaaaaataaattcaaatgatgtaaagag |
| pyrE-dndn | tttgaacgatgacctcgagcgatagccttggtcatgaagc |
| pyrE-ko-up | atcaacacactaatcggc |
| pyrE-ko-dn | taaaagcgggagaagtgc |
| pyrE-ko-ce-up | gtctctgattgatacggttg |
| IscB-ωRNA-bpr | gctcgtgttgtacaataaatgtgaggtatacgatgccaggctcttccaactttatggt |
| enIscB-ωRNA-bpr | gctcgtgttgtacaataaatgtgaggtatacgatgccaggctcgtccaactgcgg |
| HR-part-bpr-up | taatcggtatcagcacaacaaaggctcgaggtcatcgttc |
| HR-part-bpr-dn | tctttcattgtctgagagatcccctcataatttcagc |
| bpr-upup | aattatgaggggatctctcagacaatgaaagaagcggtgg |
| bpr-updn | gacggcagcttttttccactttcatcccccttttcaac |
| bpr-dnup | atgttgaaaaagggggatgaaagtggaaaaaaagctgccg |
| bpr-dndn | ttttgaacgatgacctcgagcctttgttgtgctgataccg |

| | |
|---|---|
| bpr-ko-up | gaccgtttaccttcgaagg |
| bpr-ko-dn | cggatgaataatcgcgtc |
| bpr-ko-ce-dn | gtataaagcaagctctgtg |
| enIscB-ωRNA-pps | gctcgtgttgtacaataaatgtacaccagaactcattgggctcgtccaactgcgg |
| HR-part-pps-up | ataaatctaaggcatatccggcagctcgaggtcatcgttc |
| HR-part-pps-dn | tggtttaatcctttctgagagatcccctcataatttcagc |
| pps-upup | aaattatgaggggatctctcagaaaggattaaaccacgcg |
| pps-updn | gaactggagggaatccgagcggattagcggacag |
| pps-dnup | tggcctctgtccgctaatccgctcggattccctccagttc |
| pps-dndn | cattttgaacgatgacctcgagctgccggatatgccttag |
| pps-ko-verf-up | ggcaacccaatctggatg |
| pps-ko-verf-dn | tcgaaagagagcatggaac |
| pps-ko-ce-up | caatcaaatgtgtcagcctg |
| enIscB-ωRNA-169 | gctcgtgttgtacaataaatgtaactgatgggacagctggctcgtccaactgcgg |
| HR-part-169-dn | atataaccgaccgtctgagagatcccctcataatttcagc |
| HR-part 169-up | tatttctgtcactttcacctgaagctcgaggtcatcgttc |
| 169-upup | aattatgaggggatctctcagacggtcggttatatggtcg |
| 169-updn | gggaaggagcaacaagatcaaaaaggtataaggccttaag |
| 169-dnup | cttaaggccttatacctttttgatcttgttgctccttccc |
| 169-dndn | attttgaacgatgacctcgagcttcaggtgaaagtgacag |
| 169-ko-verf-up | agatgtgaactatacgcttg |
| 169-ko-verf-dn | gattgatagcttgttactc |
| 169-ko-ce-dn | aagattacaatggagtggag |
| mcherry-up | ttacttgtacagctcgtccatg |
| mcherry-dn | atggtgagcaagggcgagg |
| mch-spo0A-part-up | cctcgcccttgctcaccatgtttcttcctccccaaatgtag |
| mch-spo0A-part-dn | catggacgagctgtacaagtaaacatgagcttattaagtggtc |
| mCherry-ce-up | gcgtgatgaacttcgagga |
| spl-up | cacaatttttgcttctcacacgtacaataccagtagatt |
| spl-dn | tttggggaggaagaaacttaaataccaaaaatgtgttgac |
| enIscB-ωRNA-rrn | gctcgtgttgtacaataaatgtcttggcagtgtggaatggctcgtccaactgcgg |
| HR-part-rrn-up | agctgggttcagaacgtcgtgaggctcgaggtcatcgttc |
| HR-part-rrn-dn | tgtaggcacacggtctgagagatcccctcataatttcagc |
| rrn-HR-upup | aattatgaggggatctctcagaccgtgtgcctacaagtag |
| rrn-HR-updn | tcaacacatttttggtatttaacaggaacttcgctactat |
| mcherry-up-rrn | tagtagcgaagttcctgttaaataccaaaaatgtgttgac |

| | |
|---|---|
| rrn-HR-dnup | cactgcgggctctttttcatggtattccacactgccaaga |
| rrn-HR-dndn | cattttgaacgatgacctcgagcctcacgacgttctgaac |
| mcherry-dn-rrn | gctttttcttggcagtgtggaataccatgaaaaagagcccg |
| rrn-ko-verf-dn | ggtcctctcgtactaaggac |
| dnaN q up | gcacttgccgcagattga |
| dnaN q dn | aatgcaagacggtggctatc |
| mcherry q up | caagctgaaggtgaccaagg |
| mcherry q dn | gtcctcgaagttcatcacgc |

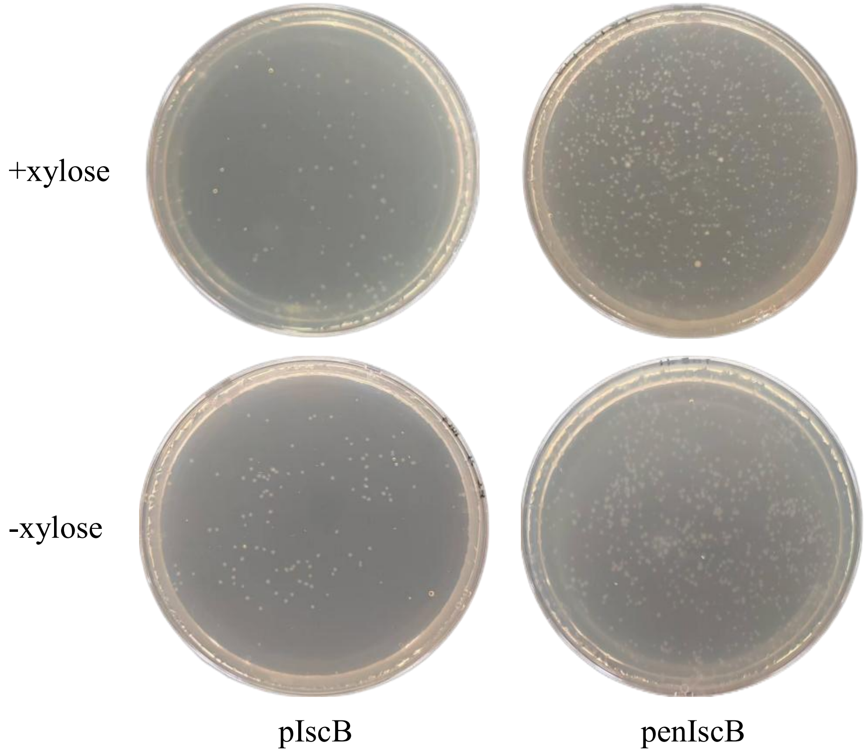


Fig S1 The toxicity tests of IscB and enIscB in *B. subtilis* SCK6. +xylose: the LB plate with 3% xylose to induce the expression of IscB/enIscB; -xylose: the LB plate without 3% xylose, and IscB/enIscB was not expressed.

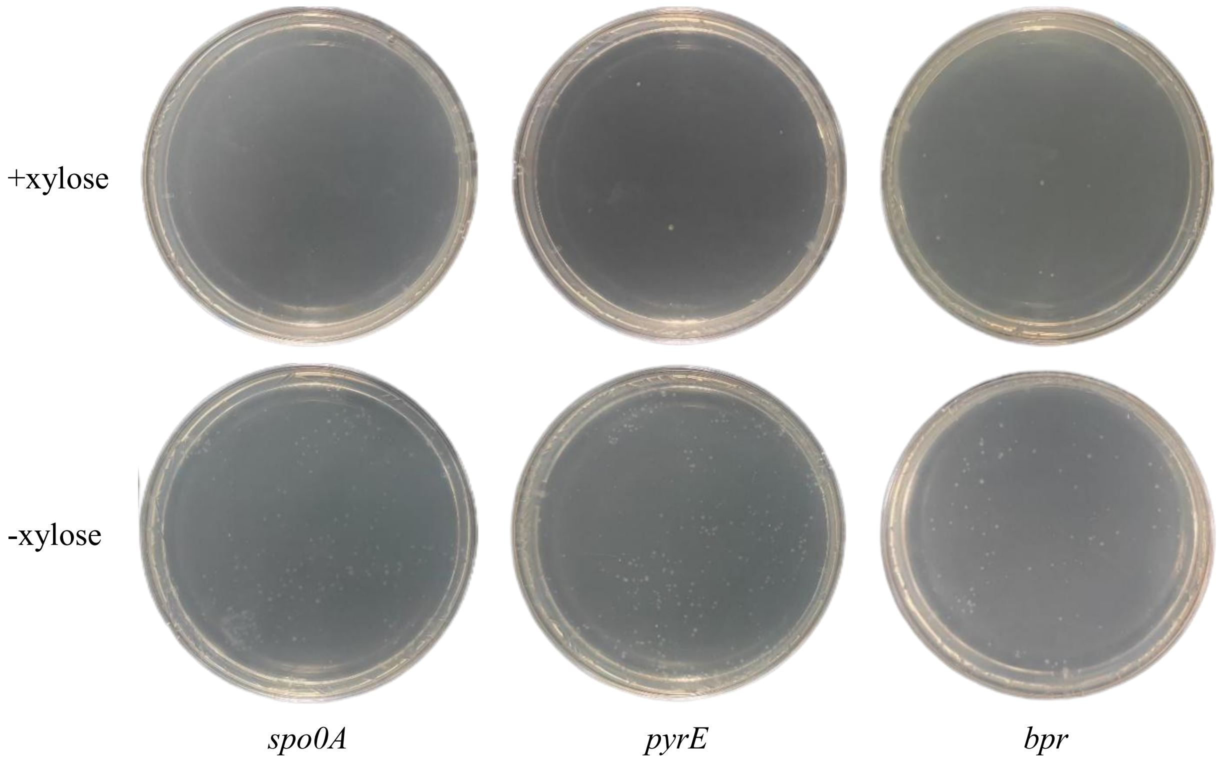


Fig S2 The test on the cleavage of the genome of *B. subtilis* SCK6 by using pBsuIscB-ωRNA. +xylose: the LB plate with 3% xylose to induce the expression of IscB; -xylose: the LB plate without 3% xylose, and IscB was not expressed.

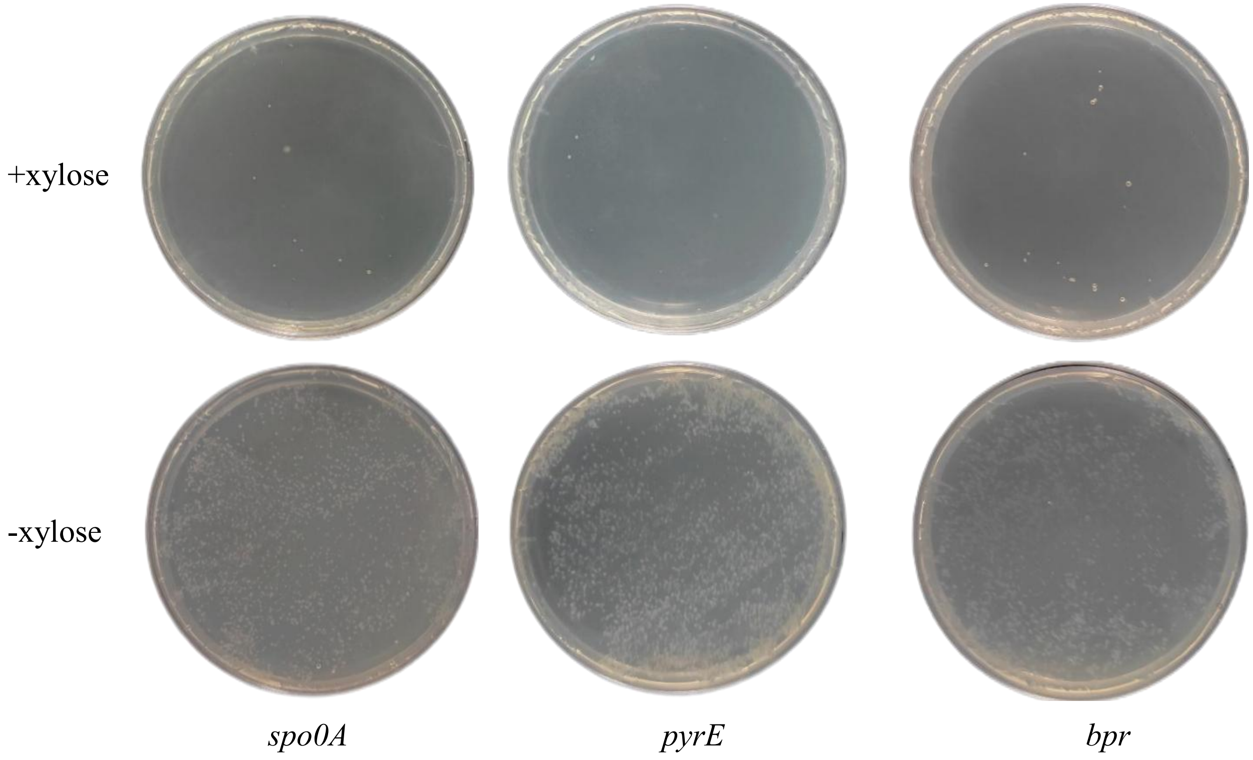


Fig S3 The test on the cleavage of the genome of *B. subtilis* SCK6 by using pBsuenIscB-ωRNA. +xylose: the LB plate with 3% xylose to induce the expression of enIscB; -xylose: the LB plate without 3% xylose, and enIscB was not expressed.

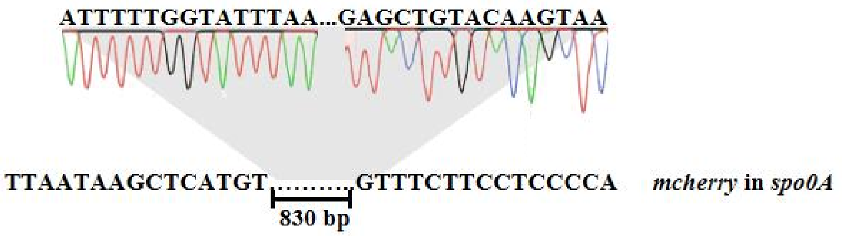


Fig S4 Sequencing results of single-copy *mCherry* gene integration at the *spo0A* site.

A

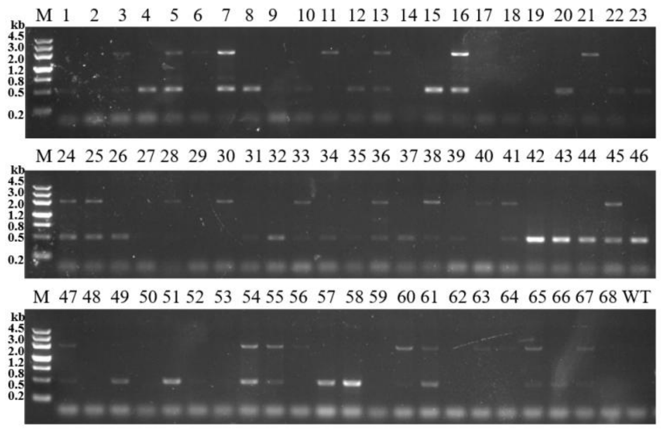


B

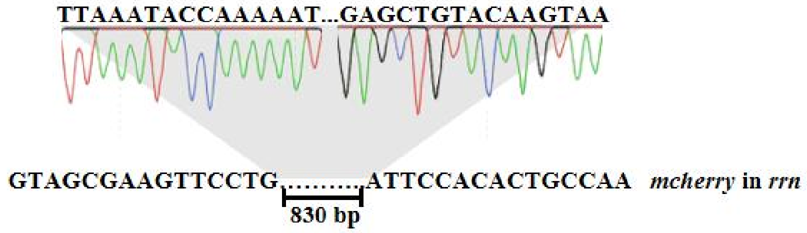


Fig S5 （A）DNA gel image of the integration of the multi-copy *mCherry* genes at the *rrn* operon. （B）Sequencing results of multi-copy *mCherry* genes integration.

REFEERENCE

(1) Yaokang, W.; Yanfeng, L.; Xueqin, L.; Jianghua, L.; Guocheng, D.; Long, L., CAMERS-B: CRISPR/Cpf1 assisted multiple-genes editing and regulation system for *Bacillus subtilis*. Biotechnol Bioeng. 2020, *117* (6), 1817-1825.
